# Aging-Associated Autoimmunity in Genetically Diverse UM-HET3 Mice Shows a Female Sex Bias

**DOI:** 10.1101/2025.07.25.666876

**Authors:** Natalia Guzniczak, Magdalena Makuch, Makayla Tillett, Harini Bagavant, Umesh S. Deshmukh

**Affiliations:** Arthritis & Clinical Immunology Program, Oklahoma Medical Research Foundation, Oklahoma City, OK 73104, USA

**Keywords:** Autoimmunity, Mice, Salivary Glands

## Abstract

C57BL/6 (B6) mice, often considered a non-autoimmune control strain, spontaneously develop autoantibodies and lymphocytic infiltration in the salivary glands (SG) with aging. However, as an inbred strain, B6 mice have a limited genetic background and do not fully represent a genetically diverse population. To assess whether genetic diversity influences the development of age-related autoimmunity, we studied UM-HET3 mice, a four-way cross that is commonly used in aging research. By 14–20 months of age, females showed significantly higher frequencies and endpoint titers of anti-nuclear antibodies. Older females also exhibited increased levels of splenic atypical/age-associated B cells and follicular helper T cells, populations associated with the production of autoantibodies. Similar immune cell changes were observed in the SG, with some female mice developing organized lymphocytic foci consisting of T and B cells. Our findings demonstrate that UM-HET3 mice naturally develop systemic autoimmunity and sialadenitis with age, with a clear female bias. Since female UM-HET3 mice have a longer median lifespan than males, these autoimmune responses may reflect benign autoimmunity, representing a heightened immune response associated with aging.

---

Sjögren’s disease (SjD) is a chronic autoimmune disorder that primarily affects the exocrine salivary and lacrimal glands (Zhao et al. 2024). It exhibits a striking female predominance and is typically diagnosed in individuals over 50 years of age (median age = 60 years, interquartile range [IQR] = 50–75 years) (Retamozo et al. 2021; Conrad et al., 2023). To explore the role of aging in SjD, we previously examined B6 mice across various age groups. We found that aged females spontaneously develop SjD, characterized by a high titer of circulating autoantibodies and well-organized lymphocytic foci in the SG (Bagavant et al. 2024). These autoimmune features are established by 14 months of age and persist through 26 months, drawing parallels with the human aging timeline for SjD onset (Flurkey et al. 2007). However, the limited genetic background of B6 mice raises the question of whether such features generalize to more genetically diverse strains. To address this, we studied autoimmune manifestations in the UM-HET3 mice, a four-way cross of B6, BALB/cByJ, C3H/HeJ, and DBA/2J strains. UM-HET3 mice are genetically heterogeneous and serve as the standard model for the National Institute on Aging’s (NIA) Intervention Testing Program (ITP) (Nadon et al., 2017).

Female and male UM-HET3 mice were obtained from The Jackson Laboratory at 3 months (n = 10/sex) and 12 months (n = 10/sex) of age. Mice were aged and euthanized at 4 months (young) and between 14 and 20 months (old). Terminal bleeds were assessed for autoantibodies using established protocols (Bagavant et al. 2024). Aged mice of both sexes developed class-switched IgG autoantibodies (Fig. 1A) with reactivity against nuclear and/or cytoplasmic antigens. Sera from young mice (n=5/sex) tested negative for autoantibodies. Endpoint titers determined by testing serial serum dilutions revealed that females had both a higher incidence and significantly elevated titers of circulating autoantibodies (Fig. 1B).

**FIGURE 1:**
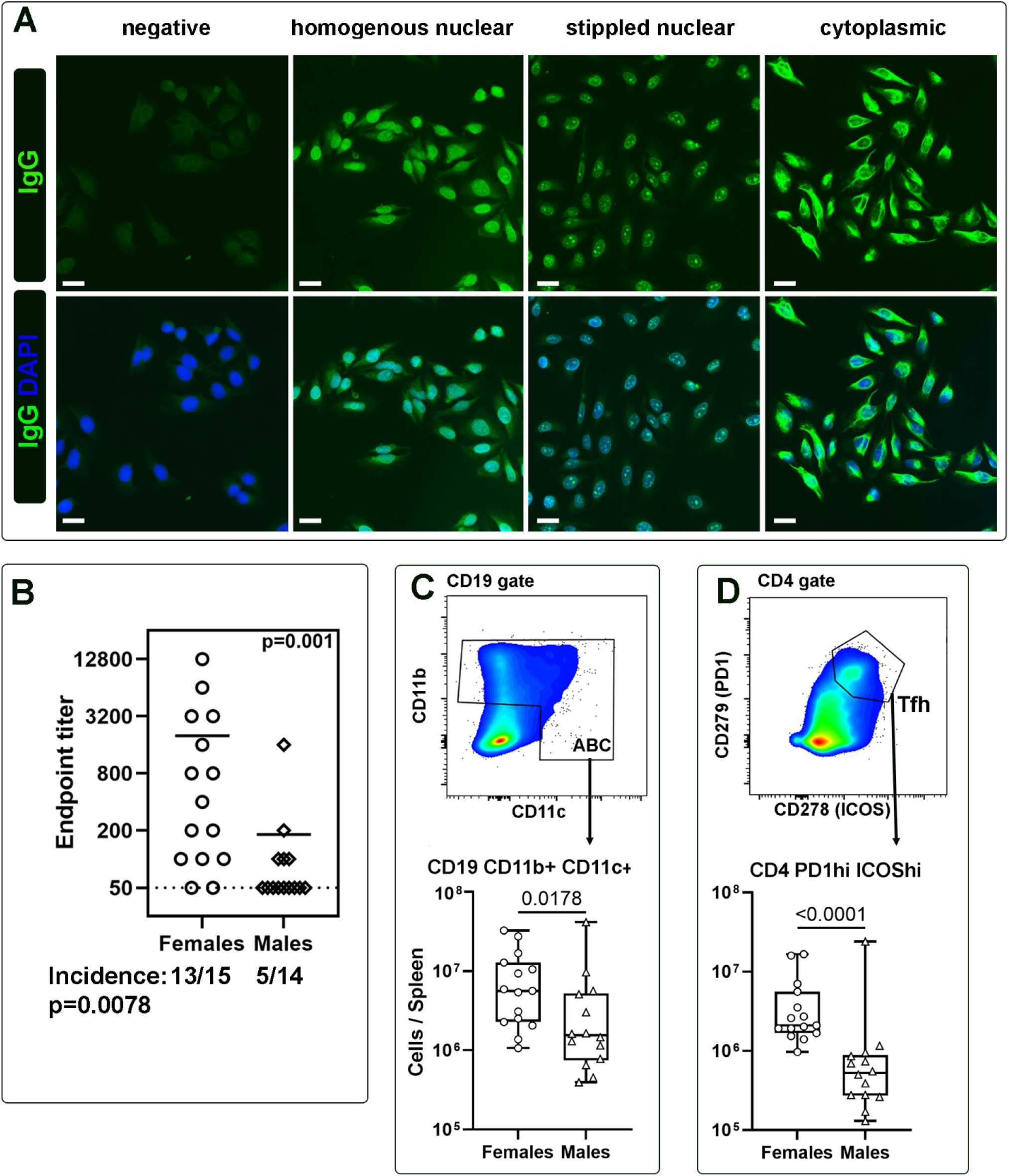
Systemic autoimmunity in 14-20-month-old UM-HET3 female (n=15) and male (n=14) mice. **(A)** IgG autoantibodies in sera of UM-HET3 mice. Young mouse sera (1:50 dilution) was used as a negative control. Representative images of different reactivity patterns: homogeneous nuclear, stippled nuclear, and cytoplasmic (scale bar:10 μm). **(B)** Endpoint titers were determined by testing serial serum dilutions. **(C)** Flow cytometry gates and numbers for atypical B cells (ABC) expressing CD11b+ and/or CD11c within the CD19+ B cell gate. **(D)** Tfh gates and numbers for cells expressing PD1 and ICOS within the CD4 T cell gate. The Mann-Whitney non-parametric test was used to calculate p-values and determine differences between the two groups.

To determine the mechanisms underlying this female-biased autoantibody response, we analyzed splenic immune cells using spectral flow cytometry (Supplemental material). The total counts of CD45+ immune cells, CD4+ and CD8+ T cells, and CD19+B220+ B cells were comparable between age-matched males and females (Table S1). However, females had significant increases in age-associated/atypical B cells (ABC), which express myeloid cell markers like CD11b and CD11c (Rubtsova et al. 2017) (Fig. 1C). Furthermore, the ABC showed a strong positive correlation with autoantibody titers (Spearman ρ=0.682, p=4.5 × 10^−5^, n=29).

IL-21 producing CD4+ T follicular helper (Tfh) cells, that support ABC differentiation, were significantly increased in females (Fig. 1D) and correlated with ABC numbers (Spearman ρ=0.732, p=6 × 10^−5^, n=29). In addition, females also showed elevated levels of CD4 effector/effector memory (CD44 hi CD62L lo) and central memory (CD44 hi, CD62L hi) subsets, indicating a global activation of CD4 T cells. However, CD8+ T cell subsets did not show significant sex differences (Table S1).

We next examined whether immune cell changes in the spleen were reflected in the SG. Significant increases in total CD45+, CD4+ T, and CD19+ B cells, as well as ABC and CD4 Tfh were noted in female SG (Fig. 2A). Also elevated were the activated CD4+ T cell subsets; CD44 hi CD62L lo effector memory, CD44 hi CD62L hi central memory, and CD44 lo CD62L lo activated effector cells (Table S1). Collectively, these data demonstrate an age-related loss of immunological tolerance and a corresponding increase in adaptive immune cell activation in female UM-HET3 mice, both systemically and in the SG.

**FIGURE 2:**
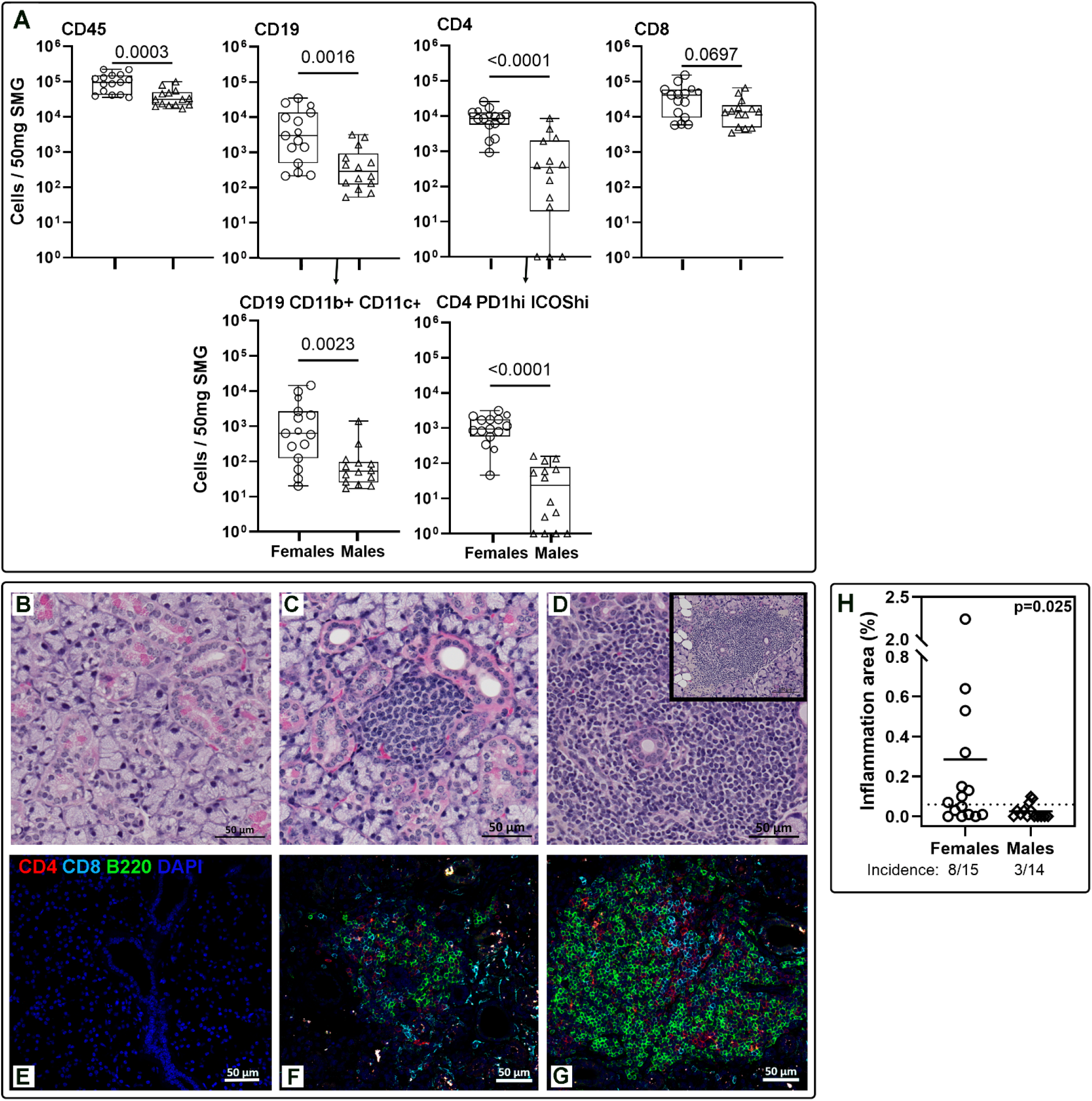
SG immune cell infiltration and lymphocytic foci in 14-20-month-old UM-HET3 female (n=15) and male (n=14) mice. **(A)** Female SGs have significantly higher numbers of immune cells (CD45+), specifically CD19+ B (including ABC) and CD4+ T (including Tfh). **(B-D)** Representative photomicrographs of Hematoxylin and Eosin-stained female SG showing no focus **(B)**, small **(C)**, and large focus **(D)**. Scale bar:50 μm. The inset **(D)** shows the entire large focus at lower magnification. Scale bar:100 μm. **(E-G)** Immunofluorescence staining for CD4, CD8, and B220 cells in SG with no focus **(E)**, small **(F)**, and large **(G)** foci. **(H)** Females show higher severity of sialoadenitis represented as the area of inflammation (the size of lymphocytic foci/ total area of the SG section) x 100. Statistical significance was determined by the Mann-Whitney test.

The mechanism by which immunological tolerance is abrogated in female UM-HET3 mice remains unclear. A recent study analyzing the liver transcriptomes of UM-HET3 mice found that cellular responses to IFN-β were more robust in females (Bou Sleiman et al. 2022). Although elevated type I IFNs are implicated in the pathogenesis of autoimmune diseases such as SjD and lupus, the IFN response in UM-HET3 mice may not reach pathogenic thresholds but may be sufficient to lower the threshold of B cell activation (Braun et al. 2002).

To determine whether these immune cell changes led to the formation of lymphocytic foci, characteristic of SjD, we examined SG histology. Sialadenitis in UM-HET3 mice exhibited a range of severity, from no inflammation to small and large foci (Fig. 2B–2D). These inflammatory foci contained CD4+ T cells, CD8+ T cells, and B cells, with B cell-rich aggregates predominating in larger foci (Fig. 2E–2G). The lymphocytic foci were more common and severe in aged females compared to males (Fig. 2H). Despite similarities in immune cell infiltrates with those of B6 mice, only a few UM-HET3 females developed severe sialadenitis comparable to that of age-matched B6 females, reinforcing the idea that genetic susceptibility is crucial for end-organ involvement in autoimmune disease. Additionally, our results suggest that in UM-HET3 mice, the genetic contributions from the three other strains might prevent the development of full-blown SjD-like autoimmunity seen in aged B6 mice.

To our knowledge, this is the first study to report systemic autoantibody responses in UM-HET3 mice. Despite the high incidence and titers of autoantibodies in aged females, the existing literature consistently shows that female UM-HET3 mice have a longer median lifespan than males (Cheng et al. 2019). This observation suggests that autoimmune responses in UM-HET3 females may reflect a benign, age-associated increase in immune activity rather than overt pathological autoimmunity. One limitation of our study is that our data comes from a single cohort of mice. Since NIA ITP studies are conducted at four independent sites, future analyses of sera from these sites could help determine whether environmental influences affect autoimmune responses in UM-HET3 mice.

Currently, more than 20 compounds are under investigation in the ITP (National Institute on Aging). It would be of considerable interest to evaluate whether any of these interventions influence autoimmunity. Such data would provide valuable insights into their immunomodulatory potential and be beneficial for potential use in preclinical models of various autoimmune disorders.

## Acknowledgments

The authors thank the OMRF Imaging Core and Flow Cytometry Core (supported by NIH Grant number: 1S10OD028479-01) for assistance.

## Authors’ Contributions

**NG, MM, MT, HB, USD**: performed experiments, analysed data, and edited the manuscript. **HB, USD**: conceived the idea, designed experiments, and wrote the manuscript.

