## Supporting Information for "Aging-Associated Autoimmunity in Genetically Diverse UM-HET3 Mice Shows a Female Sex Bias"

**Supplementary Information**

Experimental Procedures.

Female and male UM-HET3 mice were obtained from The Jackson Laboratory at 3 months (n = 10/sex) and 12 months (n = 10/sex) of age. Mice were aged and euthanized at 4 months (young, n=5 mice/ sex) and between 14–20 months (old, females n=15, males n=14). One male mouse was found dead, possibly due to fight-related injuries. Terminal bleeds were collected via cardiac puncture, and mice were perfused with PBS before tissue harvest. Submandibular salivary glands were cut into pieces and fixed in 10% neutral buffered formalin for histopathology, or 1% Paraformaldehyde-lysine-periodate for immunostaining.

For the analysis of immune cells by flow cytometry, submandibular gland pieces were weighed (56.38 ± 17.68 mg, mean ± SD, n = 29), minced, and digested with collagenase and DNAse to obtain single-cell suspensions. The cells were stained with two different antibody cocktails for T cell (CD45 BUV737, CD4 BUV563, CD8 BUV395, CD44 AF488, CD62L BV785, CD278 (ICOS) PE-Cy5, CD279 (PD1) PE-Dazzle594) and B /myeloid cell (CD45 BUV737, B220 FITC, CD19 BUV661, CD21 APC-Fire750, CD23 PE-Dazzle594, CD11c BV711, CD11b PE-Cy5, F4/80 AF647, MHC II AF700) subsets. Zombie Aqua viability kit (Biolegend) was used for live-dead cell discrimination. An average of 2.4 x $10^6$ events/sample were acquired on a 5-laser Cytek Aurora flow cytometer. The data were analyzed using FlowJo software, and the results are presented as the number of cells per 50 mg of salivary gland tissue.

Spleens were harvested, and cell suspensions were prepared using standard protocols. RBCs were lysed, and the numbers of cells per spleen were counted using a hemocytometer. Each antibody cocktail was used to stain ~1.5 million cells, and 0.5 - 1 x $10^6$ events were acquired per sample. Data are presented as the number of each cell type per spleen.

The reagents and methodologies for serum autoantibody analyses, quantification of salivary gland histopathology, immunofluorescence staining, and flow cytometry have been previously described. (Bagavant et al. 2024, Bagavant and Deshmukh 2020).

**Supplementary Table S1**: Immune cells in spleen and salivary gland of aged UM-HET3 female (n=15) and male (n=14) mice.

| | | Spleen[#] | | | Salivary Glands[$] | | |
|---|---|---|---|---|---|---|---|
| **Cell type** | **Marker** | **Female** | **Male** | **p value[@]** | **Female** | **Male** | **p value[@]** |
| Immune cells | CD45 | 1.25 ± 0.2 x $10^8$ | 1.05 ± 0.2 x $10^8$ | 0.234 | 10.6 ± 1.6 x $10^4$ | 3.9 ± 0.7 x $10^4$ | **0.0003** |
| **T helper** | CD4 | 18.5 ± 2.8 x $10^6$ | 16 ± 3 x $10^6$ | 0.354 | 91 ± 14 x $10^2$ | 13.7 ± 6.4 x $10^2$ | **<0.0001** |
| Naïve | CD44 lo CD62L hi | 6.03 ± 0.6 x $10^6$ | 7.37 ± 0.9 x $10^6$ | 0.252 | 1.38 ± 0.6 x $10^2$ | 0.28 ± 0.2 x $10^2$ | **0.0007** |
| Effector/ Effector memory | CD44 hi CD62L Lo | 8.13 ± 1.6 x $10^6$ | 5.05 ± 1.5 x $10^6$ | **0.0043** | 81.3 ± 13 x $10^2$ | 12.2 ± 5.8 x $10^2$ | **<0.0001** |
| Central memory | CD44 hi CD62L hi | 1.5 ± 0.3 x $10^6$ | 0.91 ± 0.2 x $10^6$ | **0.0068** | 2.31 ± 0.7 x $10^2$ | 0.22 ± 0.1 x $10^2$ | **<0.0001** |
| Activated Effector | CD44 lo CD62L lo | 2.03 ± 0.6 x $10^6$ | 2.18 ± 0.9 x $10^6$ | 0.683 | 4.7 ± 1.3 x $10^2$ | 0.86 ± 0.5 x $10^2$ | **0.0004** |
| Follicular helper | PD1 hi ICOS hi | 4.51 ± 1.3 x $10^6$ | 2.22 ± 1.7 x $10^6$ | **<0.0001** | 12.1 ± 2.3 x $10^2$ | 0.49 ± 0.2 x $10^2$ | **<0.0001** |
| **T cytotoxic** | CD8 | 11.1 ± 1.7 x $10^6$ | 9.57 ± 1.2 x $10^6$ | 0.621 | 4.27 ± 1 x $10^4$ | 1.85 ± 0.5 x $10^4$ | 0.0697 |
| Naïve | CD44 lo CD62L hi | 4.46 ± 0.4 x $10^6$ | 4.94 ± 0.6 x $10^6$ | 0.652 | 0.89 ± 0.3 x $10^2$ | 0.3 ± 0.2 x $10^2$ | **0.0032** |
| Effector/ Effector memory | CD44 hi CD62L Lo | 0.74 ± 0.1 x $10^6$ | 0.77 ± 0.1 x $10^6$ | 0.915 | 3.38 ± 0.8 x $10^4$ | 1.55 ± 0.5 x $10^4$ | 0.0568 |
| Central memory | CD44 hi CD62L hi | 5.05 ± 1.2 x $10^6$ | 3.3 ± 0.7 x $10^6$ | 0.234 | 22.4 ± 8.2 x $10^2$ | 1.55 ± 0.5 x $10^2$ | **0.0006** |
| Activated Effector | CD44 lo CD62L lo | 0.63 ± 0.3 x $10^6$ | 0.35 ± 0.2 x $10^6$ | 0.146 | 54.1 ± 17 x $10^2$ | 24.5 ± 6.1 x $10^2$ | 0.5045 |
| **B cell** | CD19+ B220+ | 78.6 ± 12 x $10^6$ | 62.2 ± 13 x $10^6$ | 0.201 | 8.36 ± 2.8 x $10^3$ | 0.74 ± 0.3 x $10^3$ | **0.0016** |
| Follicular | CD21 int CD23 + | 43.8 ± 5.8 x $10^6$ | 42.6 ± 8 x $10^6$ | 0.747 | 12.6 ± 6.3 x $10^2$ | 0.25 ± 0.1 x $10^2$ | **0.0006** |
| Marginal Zone | CD21 hi CD23 neg | 7.48 ± 0.9 x $10^6$ | 6.96 ± 1.7 x $10^6$ | 0.270 | 1.47 ± 0.8 x $10^2$ | 0.48 ± 0.4 x $10^2$ | 0.195 |
| Transitional | CD21 int CD23 neg | 11.3 ± 3.3 x $10^6$ | 3.6 ± 1.7 x $10^6$ | **0.0023** | 43.6 ± 14 x $10^2$ | 4.42 ± 1.5 x $10^2$ | **0.0027** |
| Double negative | CD21 neg CD23 neg | 5.75 ± 0.7 x $10^6$ | 3.87 ± 1.3 x $10^6$ | **0.0016** | 24.9 ± 11 x $10^2$ | 1.91 ± 1.1 x $10^2$ | **0.0005** |
| Atypical | CD11b/c + | 9.27 ± 2.5 x $10^6$ | 5.32 ± 2.9 x $10^6$ | **0.0178** | 26.4 ± 11 x $10^2$ | 1.65 ± 1 x $10^2$ | **0.0023** |

[#] mean± SEM cells/spleen; [$]mean ± SEM cells/50mg of salivary gland; [@] p-values calculated using the Mann-Whitney non-parametric test
